# Cancer control is a key functionality underlying evolution of extended lifespan in mammals

**DOI:** 10.1101/615914

**Authors:** Amanda Kowalczyk, Raghavendran Partha, Nathan Clark, Maria Chikina

## Abstract

The biological origin of life expectancy remains a fundamental and unanswered scientific question with important ramifications for human health, especially as the bulk of burden of human healthcare shifts from infectious to age-related diseases. The striking variability in life-span among animals occupying similar ecological niches^1^ and the numerous mutations that have been shown to increase lifespan in model organisms^2–5^ point to a considerable genetic contribution. Using mammalian comparative genomics, we correlate lifespan phenotypes with relative evolutionary rates, a measure of evolutionary selective pressure^6^. Our analysis demonstrates that many genes and pathways are under increased evolutionary constraint in both Long-Lived Large-bodied mammals (3L) and mammals Exceptionally Long-Lived given their size (ELL), suggesting that these genes and pathways contribute to the maintenance of both traits. For 3L species, we find strong evolutionary constraint on multiple pathways involved in controlling carcinogenesis, including cell cycle, apoptosis, and immune pathways. These findings provide additional perspective on the well-known Peto’s Paradox that large animals with large numbers of cells do not get cancer at higher rates than smaller animals with fewer cells^7^. For the ELL phenotype, our analysis strongly implicates pathways involved in DNA repair, further supporting the importance of DNA repair processes in aging^8–12^. Moreover, these correlations with lifespan phenotypes are consistent across the entire mammalian phylogeny, suggesting that additional constraint on these pathways is a universal requirement for long lifespan.

## Main Text

Lifespan varies dramatically (>100-fold) across mammals^1^, and numerous studies have investigated the genomic features of mammals with extreme lifespan such as bats^13,14^, naked mole-rats^15^, whales^16^, and elephants^17^. However, these studies focus on individual species that differ from their nearest sequenced relatives in millions of nucleotides. Thus, while these studies and model organism genetic studies have yielded credible candidates for genes associated with increased lifespan, it is difficult to know to what extent these represent insights into the universal mechanisms of lifespan regulation rather than species-specific adaptation or coincidental neutral changes.

The wide range of lifespans across the mammalian phylogeny (Fig. 1A) provides the ideal dataset to investigate the convergent evolution of extended lifespan from a comparative genomics context to find evolutionary commonalities underlying longevity throughout mammals. Because independent changes in lifespan occurred repeatedly in the mammalian species tree, lifespan can be viewed as a convergent trait. Molecular features that correlate with convergent changes in lifespan therefore may also occur repeatedly across a variety of organisms. In our study we use protein evolutionary rates quantified as the number of amino acid substitutions on a phylogenetic branch to infer convergent rate shifts associated with lifespan traits across the mammalian phylogeny.

**Figure 1.**
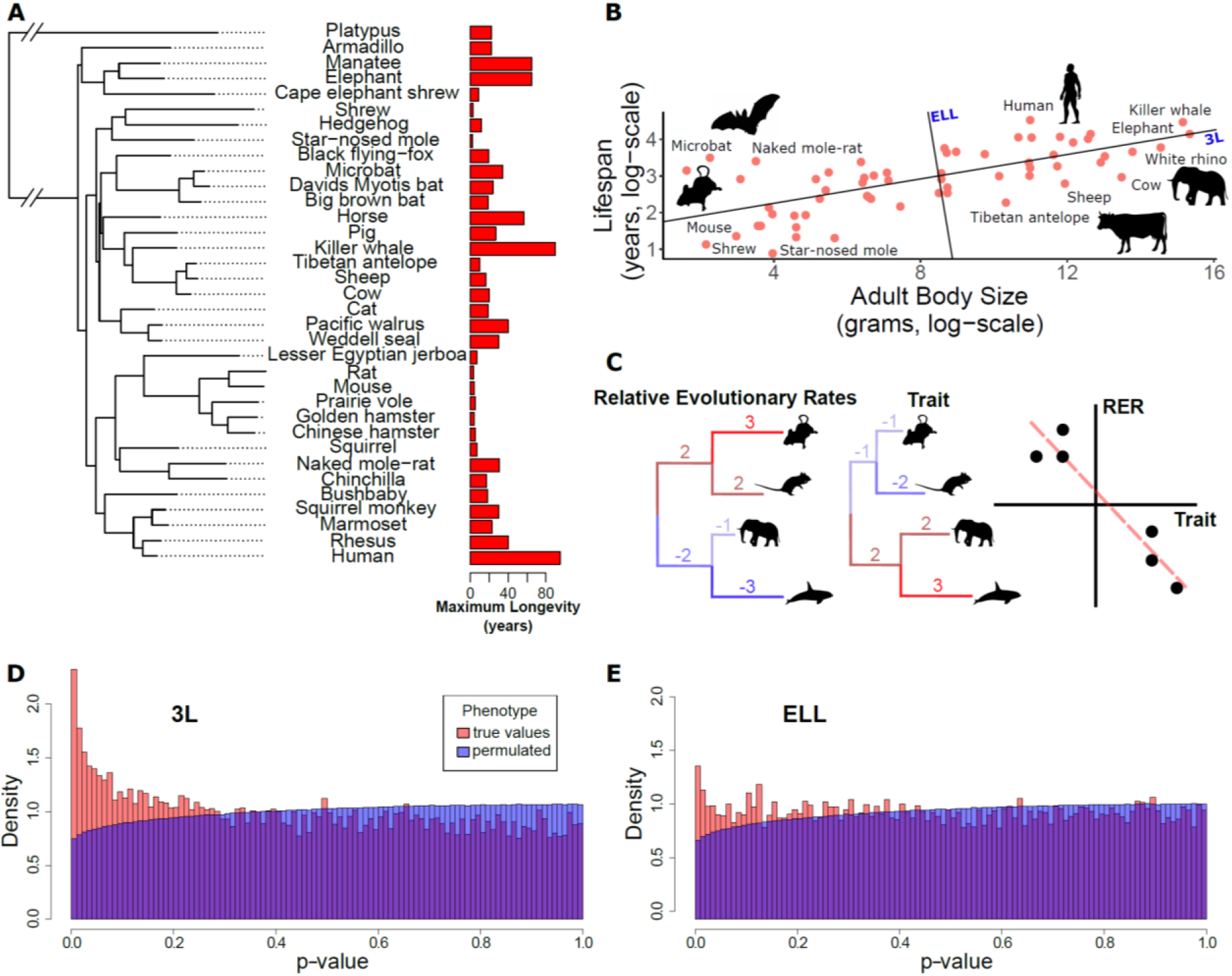
Lifespan varies widely across mammals independent of phylogeny. A) A subset of species used for this analysis alongside their maximum longevity values. B) The relationship between mammal body size and maximum lifespan. Lines represent the 3L phenotype (the first principal component between body size and lifespan) and the ELL phenotype (the second principal component between body size and lifespan). C) RERconverge pipeline to find correlation between relative evolutionary rates of genes and change in lifespan phenotypes. D) and E) Distribution of p-values from correlations between evolutionary rates of genes and change in the 3L and ELL phenotypes indicate an enrichment of significant correlations.

Evolutionary rates are useful for linking phenotypes to genes because they reflect evolutionary constraint experienced by a genetic element^6^. Genetic elements that support a specific trait are more constrained in species where the trait has a large contribution to fitness. Thus, in cases of phenotypic convergence, rates can be exploited to reveal important genes associated with the phenotype, such as changes to muscle and skin genes associated with the mammalian transition to a marine environment^18^ and loss of constraint of vision-related genetic elements in subterranean mammals^19,20^. Rate shifts can thus provide an evolutionary perspective on the contribution of genes, non-coding elements, or pathways to phenotypes of interest^21^. Here we report the genome-wide, pan-mammalian correlations between evolutionary rates of genes and lifespan phenotypes.

In mammals, lifespan is strongly positively correlated with adult body size such that the largest mammals (whales) are longest-lived and the smallest mammals (small rodents) are shortest lived (Fig. 1B). However, if lifespan is corrected for body size, species including bats, the naked mole-rat, and some primates are clearly exceptionally long-lived given their body sizes. Previous studies have focused on small numbers of species at both phenotypic extremes to find genetic and physiological explanations for their longevity^13–17^. We also study the evolution of both the “long-lived large-bodied” trait (3L) and the “exceptionally long-lived given body size” trait (ELL), so we use maximum lifespan and body size data^22^ to calculate 3L and ELL phenotypes (Supp. File 1) by extracting the first and second principal components of body size and maximum lifespan.

Having defined the 3L and ELL phenotypes, we compute the association between these phenotypes and protein-specific relative evolutionary rates using the RERconverge method^23,24^ (Fig. 1C). Relative evolutionary rates (RERs) quantify the deviation in evolutionary rate of a protein along a specific phylogenetic branch from proteome-wide expectations. Negative RERs indicate fewer substitutions than expected due to increased constraint. Positive RERs correspond to more substitutions than expected, which could arise due to relaxation of constraint or positive selection. To correlate RERs with lifespan phenotypes, we use phenotypic change along phylogenetic branches computed from maximum likelihood ancestral state reconstruction^25^. This transformation is equivalent to phylogenetically independent contrasts^26^ and thus removes phylogenetic dependence from the phenotype values.

After computing correlations between all proteins and the 3L and ELL phenotypes, we find an excess of low p-values (Fig. 1D and Fig. 1E), which indicates many genes have evolutionary rates correlated with lifespan phenotypes. In order to evaluate how much of the signal is due to true association with the phenotypes, we design a phylogenetically-restricted permutation strategy dubbed “permulations” (see Supp. Text 2). Using 1000 permulations, we find that our procedure gives conservative adjusted p-values compared to a uniform null distribution, with the fraction of non-null p-values (π_1_) equal to 0.153 for the 3L phenotype and 0.075 for the ELL phenotype (see Supp. Text 3). Overall, our analysis demonstrates a significant molecular signal for gene evolutionary rates correlated with lifespan phenotypes, with a considerably higher number of associations for the 3L phenotype.

Our analysis investigates both positive and negative correlations between evolutionary rates of genes and changes in lifespan phenotypes. Here we focus on the negative correlations because these indicate increased purifying selection for large values of the 3L and ELL phenotypes and thereby imply increased functional importance for long-lived species (see Supp. Text 7). Moreover, positively correlated genes showed no pathway enrichment and we find little evidence for positive selection (see Supp. Text 8).

While there is a clear excess of genes at low p-values, we focus on pathway enrichment results because they demonstrate stronger signals in the data and facilitate interpreting our results in the context of existing knowledge. We investigate enriched pathways for both 3L and ELL phenotypes using a rank-based method (Supp. File 2). After performing standard multiple-hypothesis testing corrections on the empirical p-values from permulations, there remains considerable pathway-level signal underlying the 3L and ELL traits.

Among pathways under increased constraint in 3L species, we find a striking abundance of pathways related to cancer control. Those pathways can be organized into the broad categories of “cell cycle control”, “cell death”, and “innate and adaptive immunity” and also include other cancer-related pathways for p53 regulation and telomere maintenance (Fig. 2A). Considering that these pathways are all related to the prevention of cancer, our 3L results can be naturally interpreted in the context of Peto’s Paradox^7^. The paradox reasons as follows: if all cells have a similar probability of undergoing a malignant transformation, organisms with more cells should have a greater risk of developing cancer. However, empirical cancer rates do not vary with body size^7^, which implies that larger animals harbor mechanisms to suppress cancer rates. Top 3L constrained pathways are associated with multiple cancer control mechanisms, including DNA repair, cell cycle control, cell death, and immune function (Fig. 2B and Fig. 2C). A normal cell’s transformation to malignancy involves failure of all these processes, and our analysis suggests that 3L animals are invested in the maintenance of each of their associated pathways through increased purifying selection. Furthermore, we find evidence for increased purifying selection in tumor suppressor genes, but not oncogenes (see Supp. Text 6). Based on enrichment and permulation results, we can infer that cell cycle fidelity, an early step in cancer development, is most important over evolutionary time scales for 3L species. Further, there is no evidence for enrichment of pathways associated with metastasis and angiogenesis, later steps in cancer development. This finding suggests that large, long-lived species have experience increased selective pressure to protect pathways involved in early cancer stages but not later stages, perhaps because the most severely negative fitness impacts of cancer are felt earlier in its development. Species-specific cancer control mechanisms have been identified in individual species, such as increased *TP53* copy number in elephants^17^, but we show here that such mechanisms likely bridge the entire mammalian phylogeny because top enriched pathways for the 3L phenotype do not depend on a handful of species (see Supp. Text 4, Supp. Text 5, Supp. Fig 1, and Supp. Fig. 2).

**Figure 2.**
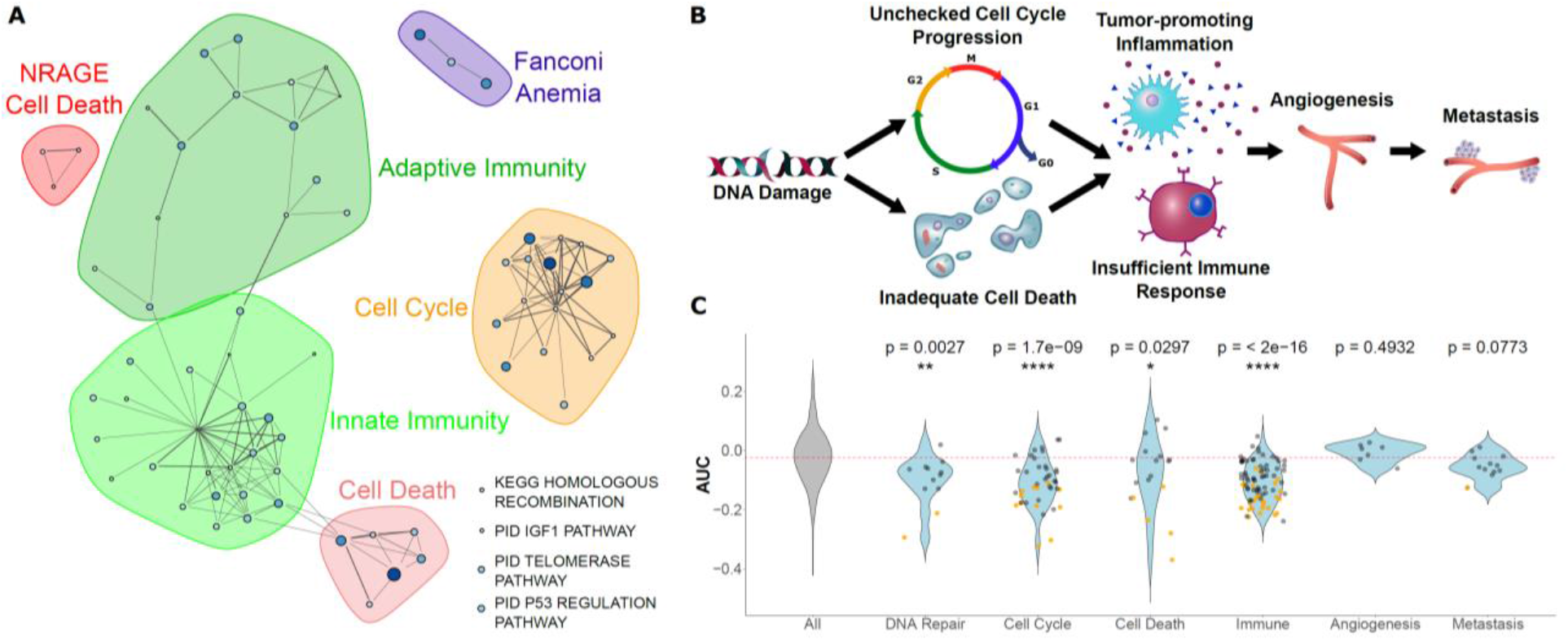
A) Significantly enriched pathways under increased constraint in species with larger values of the 3L phenotype (adjusted p-value less than 0.05, permulation p-value less than 0.05). Each dot represents a pathway, and the size and color of the dot represents the negative log of the rank-sum enrichment statistic. Width of lines connecting pathways represent the number of genes the pathways have in common. Pathways were clustered based on these connections, and shaded shapes represent this clustering. B) and C) Pathways under increased constraint in 3L species play various roles in cancer control. Pathways associated with early stages of cancer development (DNA repair, cell cycle control, cell death, and immune functions) are significantly enriched, while pathways for later stages of cancer development (angiogenesis and metastasis) are not enriched. In C), each dot represents a pathway. Yellow dots have significant permulation p-values while black dots do not. Note that dots for “All” pathways are excluded for the sake of clarity.

An additional pathway that shows a strong signal of increased constraint with the 3L phenotype is the insulin-like growth factor (IGF1) signaling pathway (Fig. 3), which deserves special consideration due to its well-established effect on lifespan in model organisms^5,27,28^. We find that IGF pathway genes evolve more slowly in large, long-lived species, suggesting that this pathway is not a likely source of molecular innovation contributing to the large and long-lived trait. This result is at odds with experimental evidence that shows that genetic manipulation of IGF signaling results in increased lifespan in diverse organisms^5,27–30^. A possible explanation is that the IGF1 pathway plays an important role in cancer control^31^, and this cancer-related signal of constraint dominates any adaptive signal related to lifespan. There are also further reasons to believe that the IGF1 pathway is not the main source of the 3L trait. Across the mammalian phylogeny, lifespan is strongly correlated with body size, but genetic perturbations in the IGF pathway result in longer-lived individuals that are of smaller size^29,32^. Furthermore, it has been shown that plasma IGF1 levels are negatively correlated with body size across diverse species of mammals^33^. Together with our findings, these observations strongly suggest that changes in the IGF1 pathway are unlikely to drive the large, long-lived phenotype, which is established through a different, yet-unknown mechanism.

**Figure 3.**
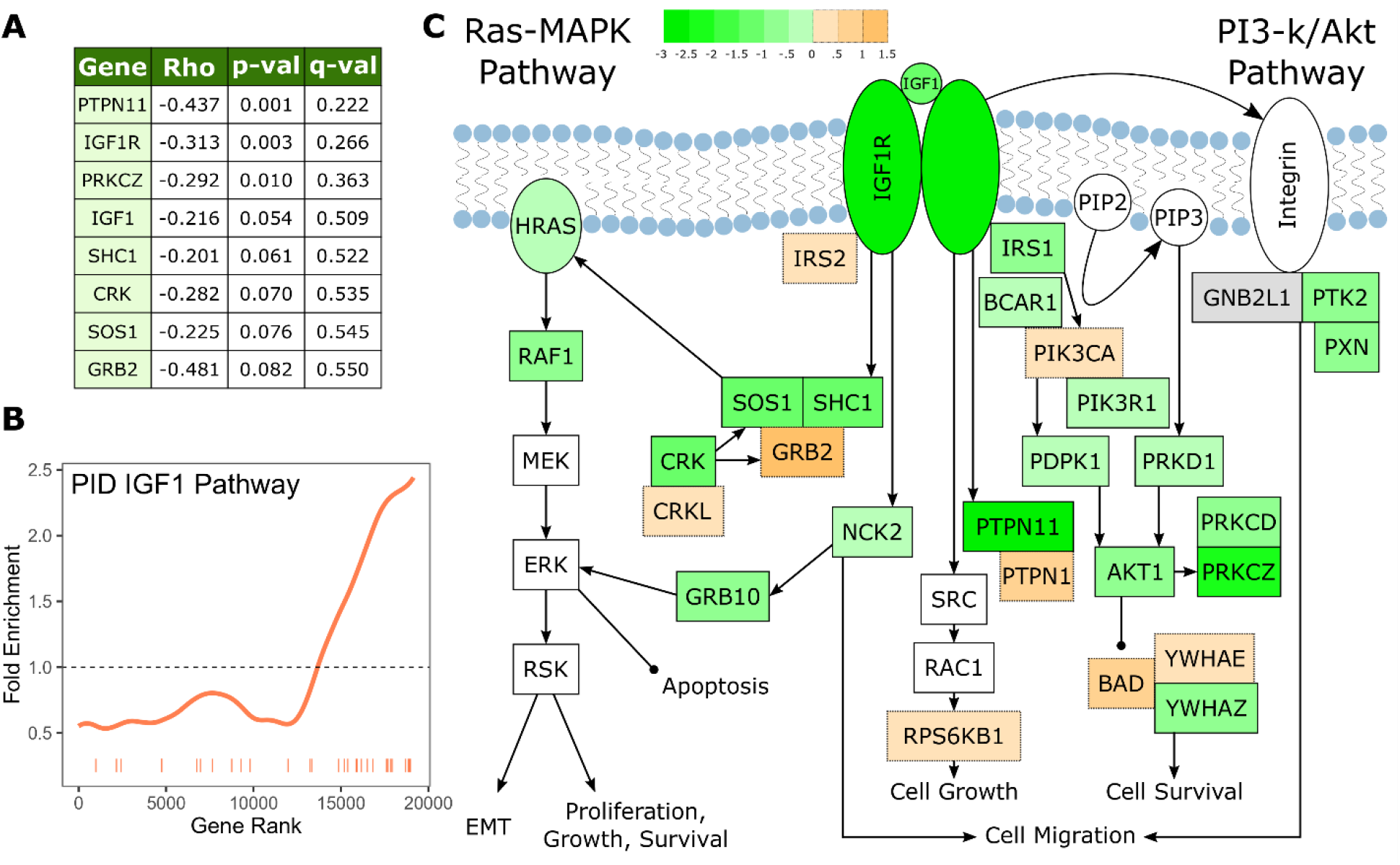
A) IGF1signaling pathway genes are significantly correlated with change in the 3L phenotype. B) The IGF1 signaling pathway is significantly enriched for increased evolutionary constraint in large, long-lived species. The barcode indicates ranks of genes in the pathway within the list of all pathway-annotated genes. The worm indicates enrichment as calculated by a tricube moving average. The dashed horizontal line marks the null value indicating no enrichment. C) The IGF1 signaling pathway contains many genes whose evolutionary rates are negatively correlated with the 3L phenotype. Shading indicates the Rho-signed negative log p-value for the correlation. Genes in white are not included in the IGF1 pathway annotation used to calculate pathway enrichment statistics, but they are included in the diagram for sake of completeness. The GNB2L1 gene (gray) is in the IGF1 pathway annotation, but correlation statistics were not calculated for that gene because too few branches in the gene tree met the minimum branch length cut-off.

For the ELL phenotype, we find a smaller, more focused set of enriched genes and pathways. Of those significantly enriched constrained pathways for ELL, we see some overlap with functional groups represented in results from the 3L phenotype, notably immune-related and DNA repair pathways (Fig. 4A). However, although the functional groups are the same, the pathways contained within them differ between the two phenotypes (Supp. File 2, Supp. File 3). In particular, the only significantly constrained DNA repair pathways for the 3L phenotype involve Fanconi’s anemia, while the ELL phenotype shows significantly constrained DNA repair pathways for a variety of repair functions (Fig. 4). Such pathways stand out not only because of the connection between DNA repair and cancer control, but also because of the observed relationship between DNA repair and aging independent of cancer incidence. This relationship can be demonstrated experimentally by creating double-stranded DNA breaks in laboratory mice to induce an aging phenotype^10^. There is also evidence that DNA damage causes dysregulation of the cellular chromatin state and thus can contribute to aging even in post-mitotic cells^34,35^.

**Figure 4.**
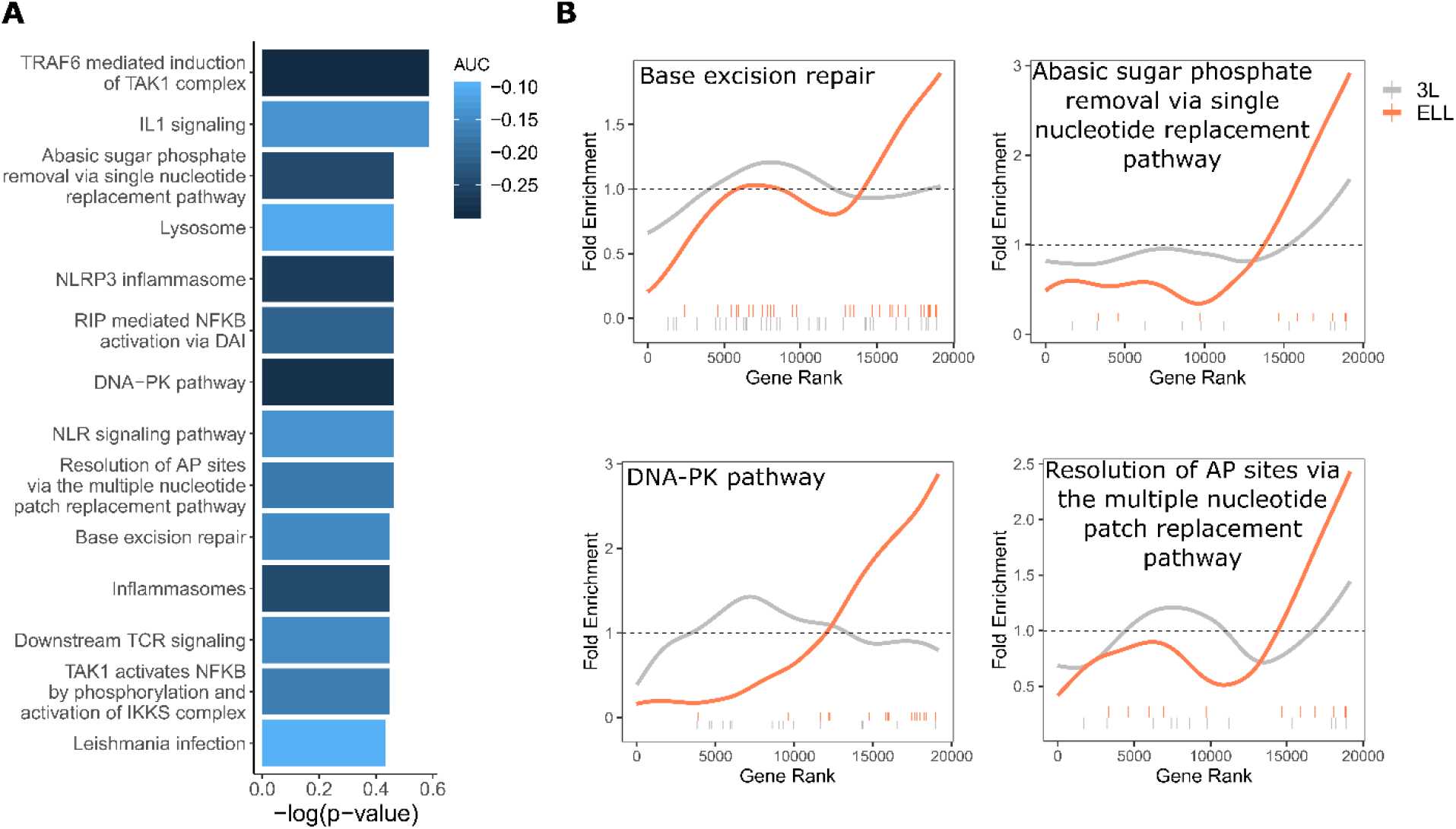
A) Significantly enriched pathways under increased constraint in species with larger values of the ELL phenotype. Bar height indicates the negative log permulation p-value for each pathway, and the color of bars indicates the pathway enrichment statistic. B) DNA repair pathways are more significantly enriched for increased evolutionary constraint in species with large values of the ELL phenotype than species with large values of the 3L phenotype. The barcodes indicate ranks of genes in the pathways within the list of all pathway-annotated genes. The worms indicate enrichment as calculated by a tricube moving average. The dashed horizontal lines mark the null value indicating no enrichment.

In addition to DNA repair-related pathways, we also noted pathways related to NFKB signaling for which overexpression in downstream targets has been associated with aging. Experimental evidence suggests a connection between NFKB signaling and DNA repair through sirtuins, a chromatin regulator family that has already been implicated in lifespan control^36,37^. Sirtuins mediate DNA damage-induced dysregulation and are also responsible for silencing NFKB-regulated genes, thus connecting the two processes^38^. Overall, our analysis strongly suggests that fidelity in DNA repair and NFKB signaling contributes to the fitness of ELL species, indicating that these pathways may be a fruitful avenue for aging research and intervention.

Together, these pathways encompass functionalities underlying the evolution of extended life, and they therefore represent candidate genes and pathways for further experimental exploration. Importantly, such genes were uncovered using an unbiased, genome-wide pan-mammalian scan. As such, these results point to keys to exceptional longevity that are not specific to one or a handful of species, but are universal across mammals.

## Acknowledgments

We would like to thank Dr. Andreas Pfenning and Dr. Dennis Kostka for helpful discussion and feedback, as well as all members of the Clark and Chikina labs.

